# Achieving Single Cell Acoustic Localisation with Deactivation Super Resolution

**DOI:** 10.1101/2024.09.20.614052

**Authors:** Cameron A. B. Smith, Mengtong Duan, Jipeng Yan, Laura Taylor, Mikhail G. Shapiro, Meng-Xing Tang

**Affiliations:** Department of Bioengineering, Imperial College London, London SW7 2AZ, UK; Division of Chemistry and Chemical Engineering, California Institute of Technology, Pasadena, California 91125, USA; Andrew and Peggy Cherng Department of Medical Engineering, California Institute of Technology, Pasadena, California 91125, USA; Howard Hughes Medical Institute, Pasadena, California 91125, USA

## Abstract

Super-resolution optical microscopy enables optical imaging of cells, molecules and other biological structures beyond the diffraction limit. However, no similar method exists to super-resolve specific cells with ultrasound. Here we introduce Deactivation Super Resolution (DSR), an ultrasound imaging method that uses the acoustic deactivation of genetically encodable contrast agents to super-resolve individual cells with ultrasound as they navigate through structures that cannot be resolved by conventional imaging methods. DSR takes advantage of gas vesicles, which are air-filled sub-micron protein particles that can be expressed in genetically engineered cells to produce ultrasound contrast. Our experimental results show that DSR can distinguish sub-wavelength microstructures that standard B-mode ultrasound images fail to resolve by super- localizing individual mammalian cells. This study provides a proof of concept for the potential of DSR to serve as a super-resolution ultrasound technique for individual cell localization, opening new horizons in the field.

---

Photo-activated localization microscopy (PALM)^1^ has been a great advance in optical imaging, allowing for the breaking of the diffraction limit by utilising activated fluorescent markers to generate super-resolved images. These principles have also been extended to the field of acoustic super-resolution^2^ which is often referred to as super resolution ultrasound (SRUS) or ultrasound localisation microscopy (ULM). Here sparse microbubbles injected into the vasculature can be imaged, localised and tracked to produce super-resolution images of the microvasculature. This technique is showing great promise as a tool for cardiovascular^3–6^, neurovascular^7,8^, and tumour imaging^9,10^.

In addition, the potential also exists for applications utilising a wider range of contrast agents, which this work hopes to explore.

Gas vesicles (GVs) are sub-micron air-filled protein-shelled particles^11^, naturally expressed by buoyant photosynthetic microbes^12^ and are capable of being expressed in bacterial and mammalian cells as acoustic reporter genes^13–15^. GVs and GV-expressing cells have been shown to be capable of extravasating out to the surrounding tissue^16^, allowing for the potential of contrast-enhanced ultrasound imaging throughout the body. Two methods are predominantly used as a means of imaging GVs in a way that segregates them from the surrounding tissue. The first is based on amplitude modulation^17–19^, which capitalises on the fact that GVs are non-linear oscillators that buckle non-destructively in an acoustic field^20,21^. The second approach capitalizes on the irreversible collapse of GVs by high-pressure pulses through a method called BURST^22^, which transmits a series of high-pressure pulses. The applied pressure is great enough that GVs are collapsed in the first high pressure pulse, such that the echoes elicited by the initial high-pressure pulse contain signals from both the GVs and the background tissue, whereas signals arising from subsequent pulses represent only the background tissue. A spectral unmixing technique is then deployed to segregate the GV signal. This effect is amplified by transmitting multiple-cycle pulses, which cause the gas released from the GVs to cavitate, producing a strong signal with enhanced contrast when compared to the subsequent frames. Sawyer *et. al*. found a linear correlation using BURST imaging between the number of bright spots on BURST imaging a beaker of liquid and the cell concentration within that liquid, suggesting that BURST imaging may be capable of detecting individual GV-expressing cells.

In this work we develop a new imaging technique based on the concepts behind BURST which is capable of imaging the majority of cells highly expressing GVs and super-localising these individual cells in a method that we call Deactivation Super Resolution (DSR) (Figure 1, Figure 2).

**FIG 1.**
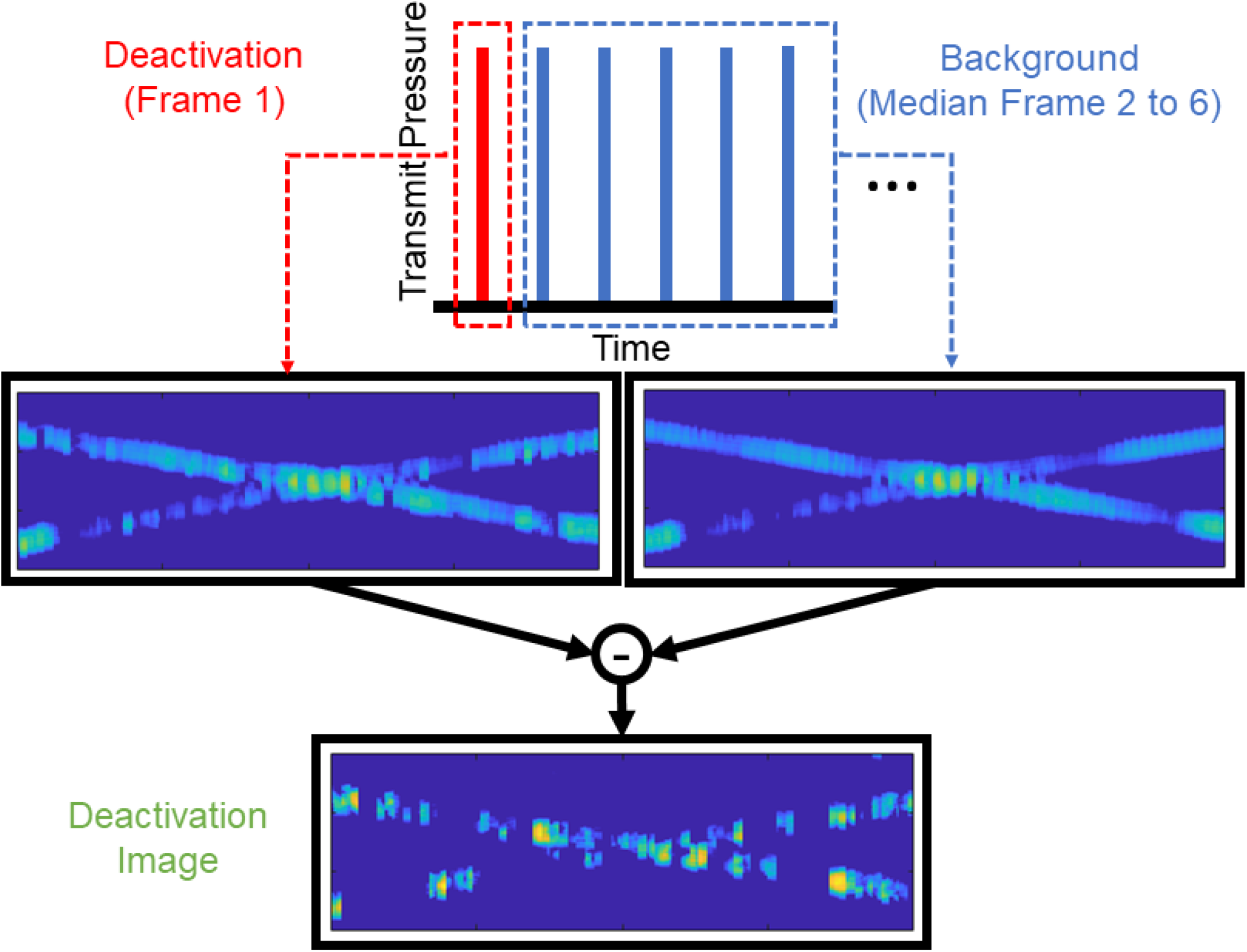
Schematic demonstrating the pulse structure and the processing involved in generating deactivation images.

**FIG 2.**
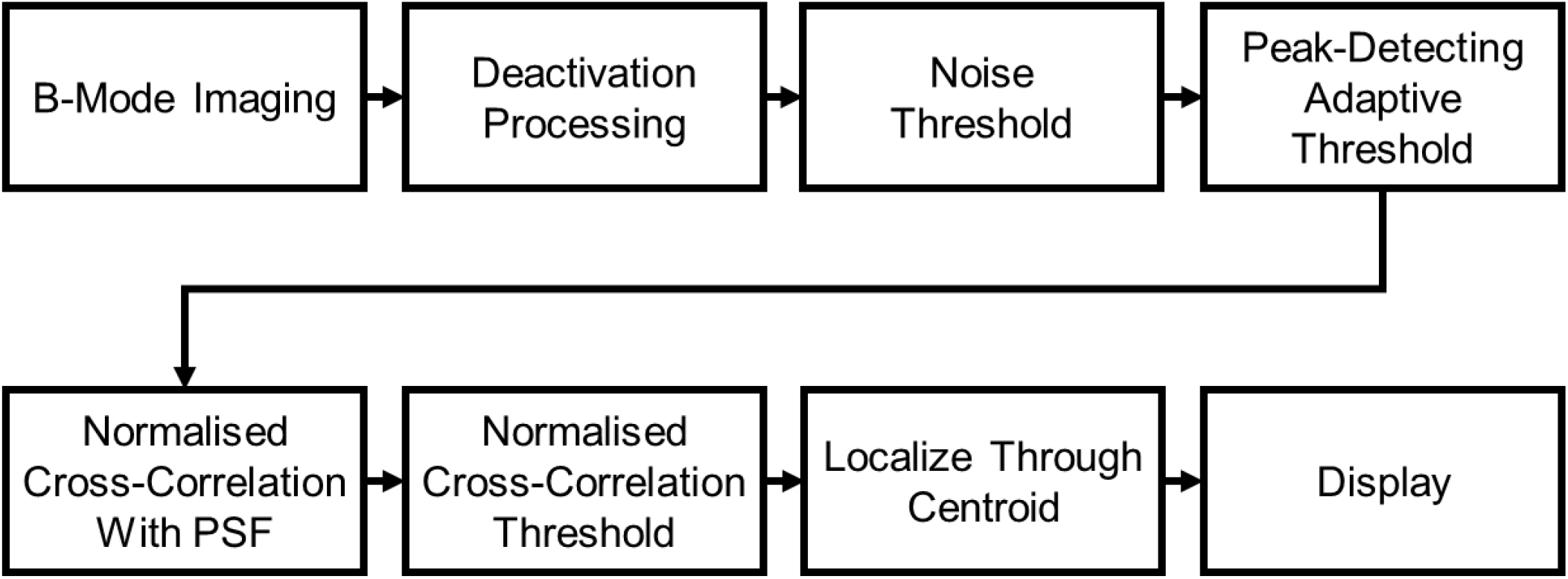
Schematic diagram showing the processing pipeline for deactivation super-resolution.

## Results

In the DSR method we use a series of six high pressure (6.2 MPa peak positive pressure) pulsed B-Mode images. In the first of these six pulses (here called the deactivation pulse) the protein shell of the GV is fractured, releasing the gas within. This gas then cavitates, releasing a strong acoustic signal, after which the bubble collapses. On the subsequent pulses the GVs are no longer present, as such the signal that is reflected to the transducers for these pulses represent all the structures in the imaging field except for the GVs. This provides us with an opportunity. By taking the median image of the five images after the deactivation pulse and subtracting this from the deactivation pulse we end up with a deactivation image which contains the isolated signal from the GVs (Fig. 1). We then use the DSR pipeline to super-localise individual cells.

To gauge the effectiveness of DSR, we used a cross tube phantom^23–25^ which comprises of two thin-walled microvessels of internal diameter 200 μm ± 15 μm in a cross formation, through which we flowed GV-expressing mammalian tumour cells^13^ with a syringe pump and imaged them with our DSR pipeline using an L11-4v probe and using 4 MHz 3 cycle pulses. The cells we used were MDA-MB-231-mARG_Ana_ cells as described in Hurt *et. al*.^13^. In brief, MDA-MB-231 cells were engineered to constitutively co-express the plasmids shown in Figure 3 (top) which enable expression of GVs derived from anabaena, GFP, and BFP when induced with the small molecule doxycycline. Images of the two crossing tubes (Figure 3) show deactivation frames showing the cells which will later be localised. Final SR images of the phantom show the flow path of cells super-localized to the centre of each channel, resulting in tighter signal distributions across the widths of each tube (Figure 4). Cross sectional signal profiles reveal that in the first location, where the tubes are further apart, the two tubes are distinguishable on both B-Mode and SR. However, in the second location, where the tubes are closer together, they are only distinguishable in SR. The average separation distance between the two tubes, defined as the closest distance where the two full width at half maximum (FWHM) profiles do not overlap is 0.83 ± 0.01 mm for the B-Mode and 0.30 ± 0.01 mm for the DSR (n = 3). The average FWHM for the B-Mode image was 0.82 ± 0.00 mm, while for the SR image it was 0.21 ± 0.01 mm (n = 6, mean ± standard error).

**FIG 3.**
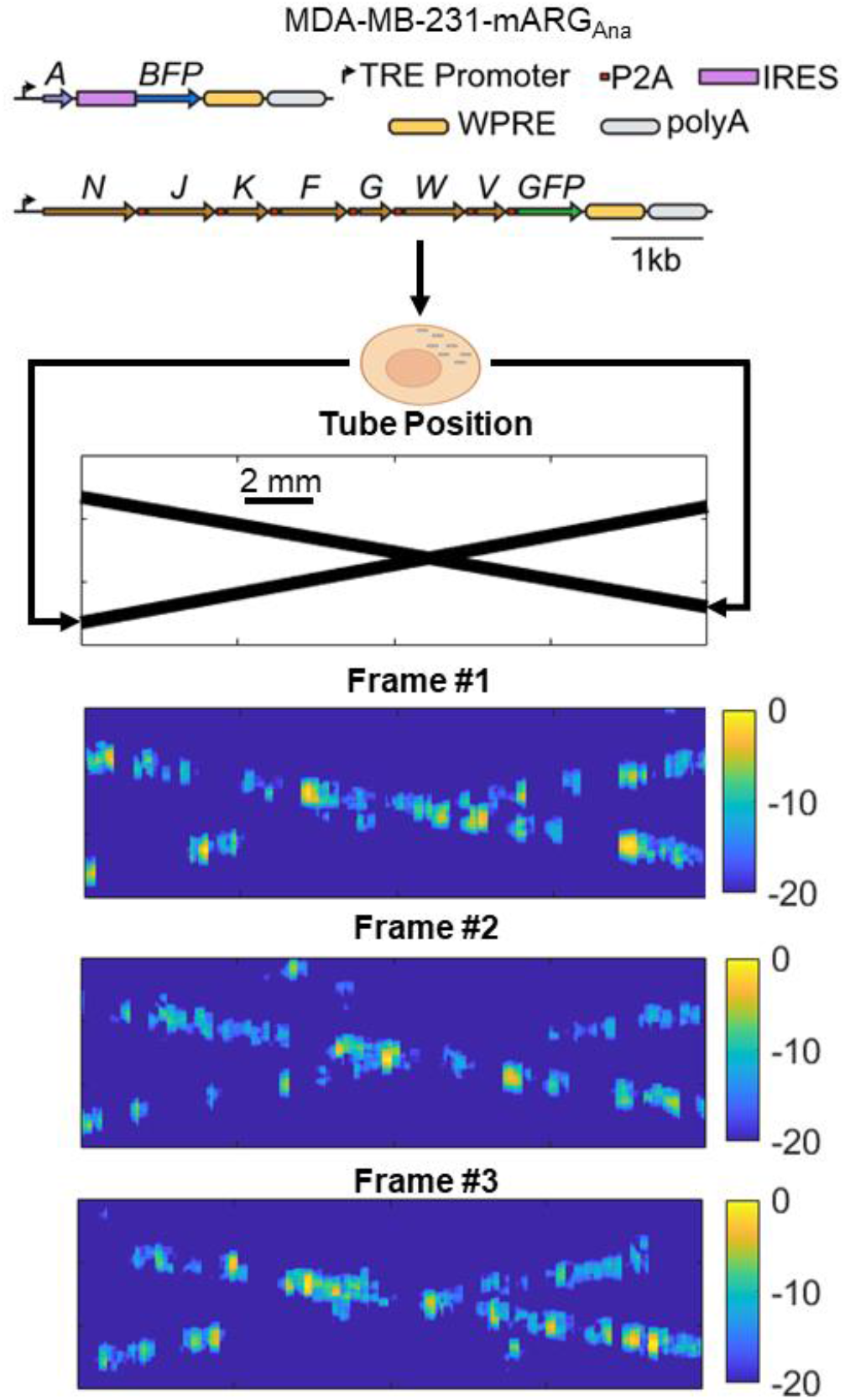
schematic diagram (top) showing the genetically engineered MDA cells (adapted from ref. ^13^) flowing through the tube in set positions, and below representative sample deactivation image frames.

**FIG 4.**
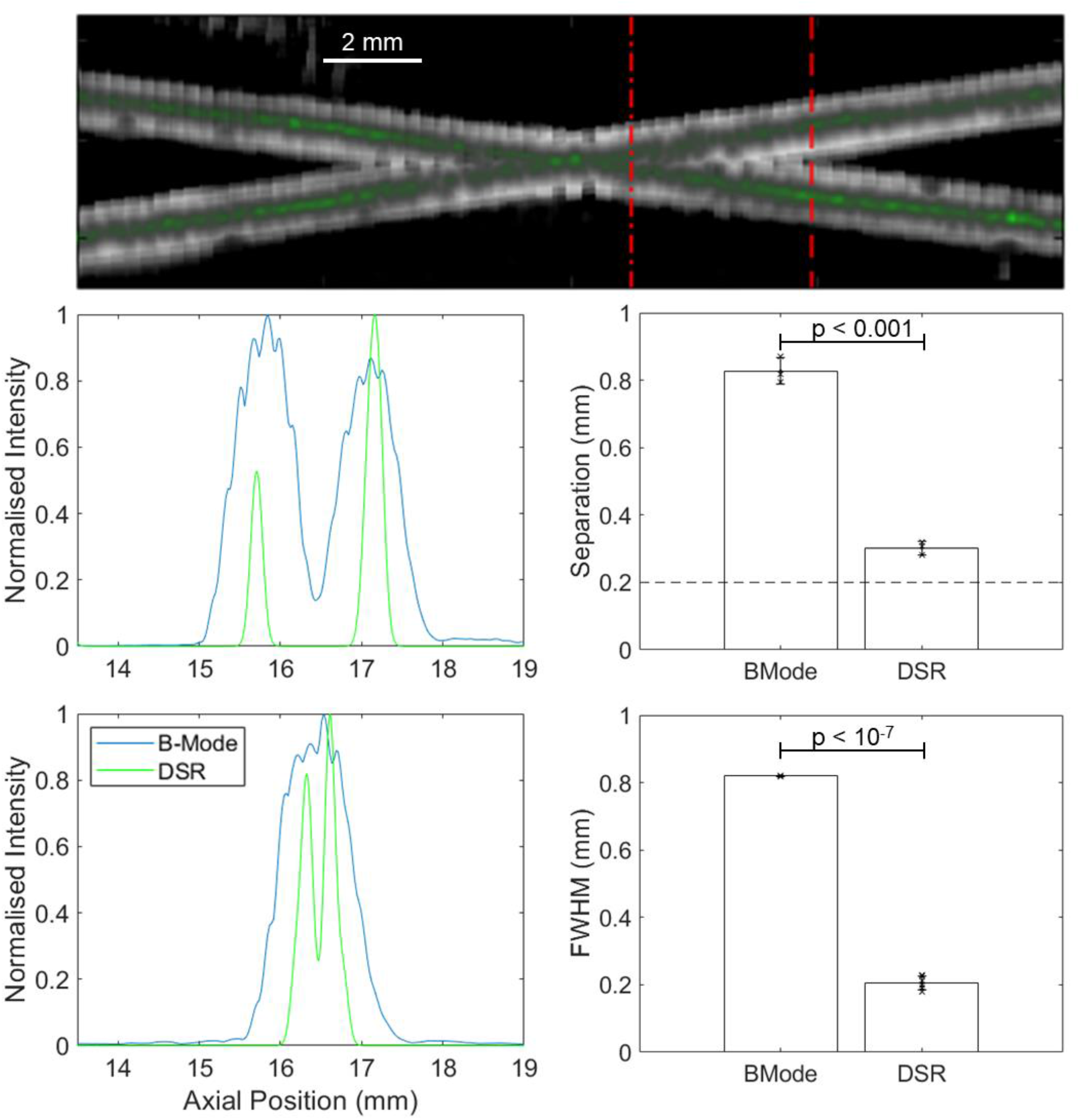
Representative deactivation super resolution image (top) and two cut through line plots showing the separation of the tubes at two positions, bar charts showing the mean and standard deviation of FWHM and tube separation distances (n = 3).

To test the sensitivity of DSR we conducted an experiment whereby we flowed cells at a variety of concentrations through the cross-tube phantom, imaging as we went. In doing so we could compare the number of cells localised against the number of cells we would expect to see based on the volume of liquid being imaged and the concentration of cells as determined by haemocytometer cell counting. We then developed a buoyancy purification technique based on centrifugating GV-expressing cells in liquids of varying density (ranging from 1.2 to 1.4 g ml^-1^). Cells that express more GVs have a lower volume-averaged density and therefore float to the top in less-dense media. After separating cells using this method, we repeated cross-tube imaging at different levels of purity (Figure 5). It was found that for the base induced cells roughly 3 % of the cells produced enough GVs to be individually localised, for the cells purified at 1.4 g ml^-1^ 29 % could be localised, and for cells purified at 1.2 g ml^-1^ this value was 81 %. Additionally, we placed the samples through a flow cytometer and investigated the proportion of cells that were expressing GFP and BFP across the different purification levels. The results of this can be seen in Figure 6. As expected, cells that were not induced with doxycycline were largely non-fluorescent in BFP and GFP, whereas the induced cells there is roughly a 78 % double positive population. In the Pure and Purer buoyancy purified cell lines we see roughly 66 % and 71 % double-positive (BFP and GFP) populations, respectively. This suggests that DSR imaging is a more effective and direct method to quantify GV expression levels in a cellular population than a fluorescence-based surrogate.

**FIG 5.**
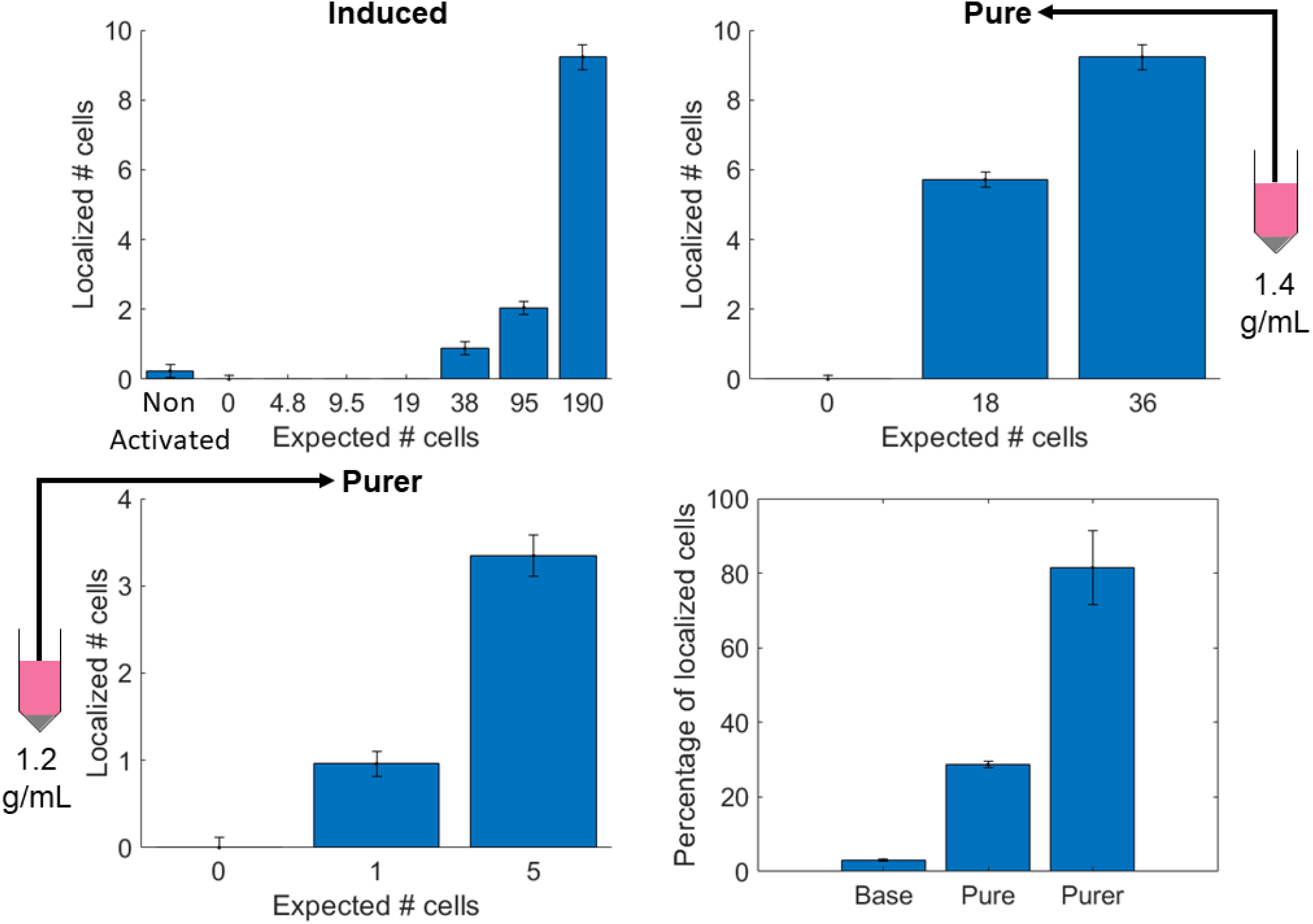
Concentration sweeps demonstrating the number of cell localisations with deactivation super-resolution imaging compared to the expected number of cells based on the cell concentration flowing at 3 levels of buoyancy purification, along with a summary figure (bottom right) showing the percentage of localised cells across the different concentrations for each purification method.

**FIG 6.**
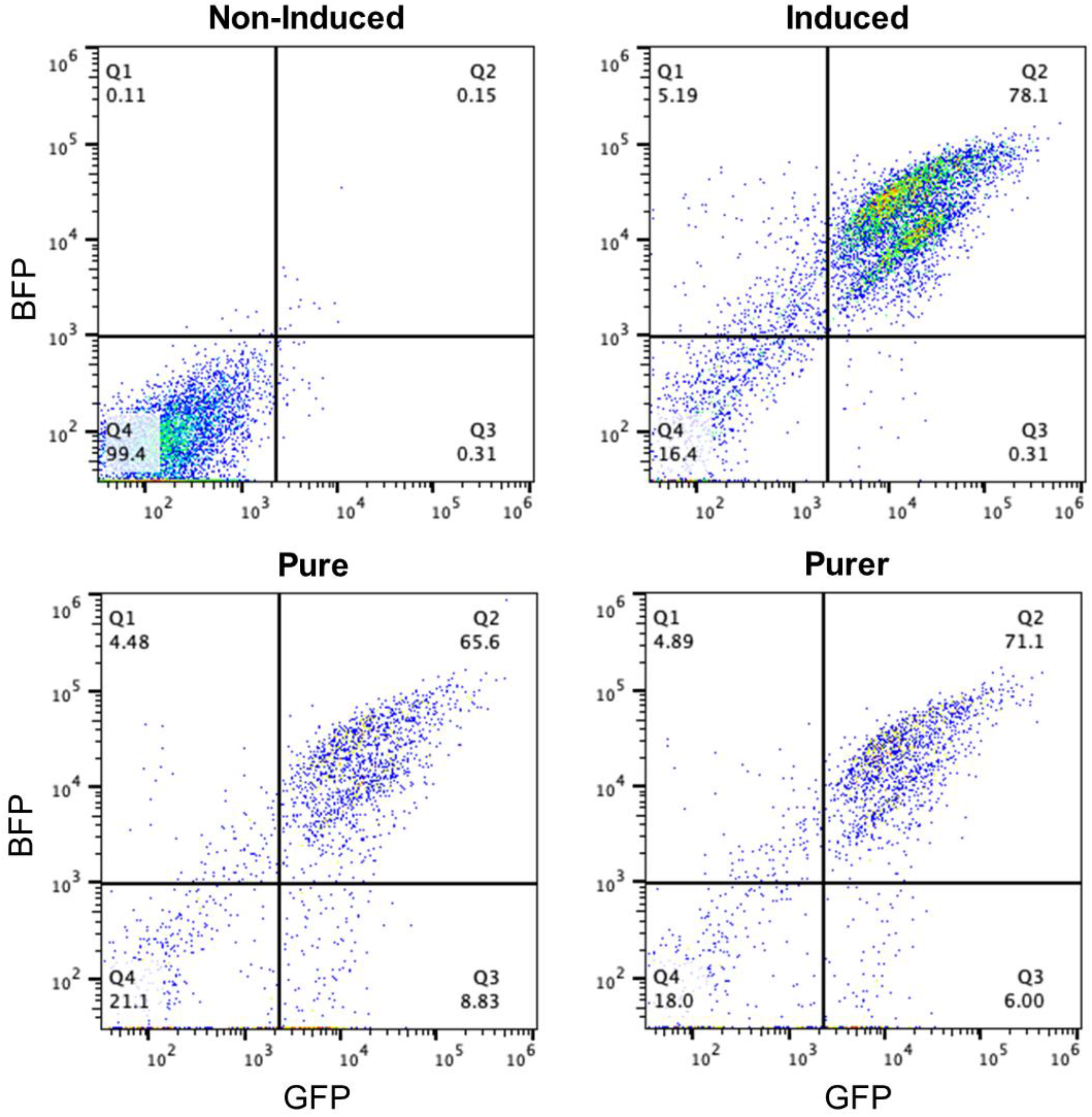
Flow cytometry showing GFP and BFP expression across induced, non-induced, and purified cells.

## Discussion

In this work we investigated the capability of the DSR technique, achieving a separation distance of 0.30 mm, which is a notable improvement over the traditional axial resolution limit without super resolution of 0.56 mm^26^. Additionally, it is notable that the FWHM and separation distances for the B-mode image are comparable, but this was not the case for the DSR method. The FWHM of the DSR method was found to be 0.21 mm, which is only marginally larger than the tube diameter of 0.2 mm, suggesting that the true FWHM and separation of the imaging technique may be smaller than this experiment suggests, and therefore perhaps that the true separation distance too is here overestimated.

Looking now to the results of the buoyancy purification experiments, it is evident that as the purification levels increase the percentage of cells that are localised relative to the number of expected cells increases dramatically, up to around 80 %. This suggests that the DSR technique is capable of imaging individual cells, and that the technique could be useful as a method to determine GV levels inside cells.

When it comes to the flow cytometry data, we find both that the percentage of cells that are GFP and BFP positive does not increase due to buoyancy purification, and also that the strength of the fluorescence is unchanged. This suggests that while GFP and BFP gating can be informative in determining whether cells are producing GVs or not, outside of this binary there is no correlation between levels of fluorescence and levels of GV expression. This also implies that buoyancy filtration cannot be substituted by fluorescence-based means, and that fluorescence-based estimates of GV expression are insufficient, highlighting the importance of acoustic-based methods such as the one proposed. Genetic engineering efforts are ongoing to ensure more consistent expression of GVs across a cell population^27^.

In conclusion, we have demonstrated that DSR is an effective super-resolution technique *in vitro*, producing images with an approximately 4-fold improvement in resolution compared to conventional ultrasound imaging. DSR has proven to be capable of localizing individual GV-expressing mammalian cells. Future studies are needed to demonstrate how this capability can be used super-resolve biological phenomena *in vivo* and beyond the vascular space in basic biology and potential clinical applications. In the meantime, in vitro cross-tube imaging provides a useful method to assess GV expression levels inside cells and evaluate the achievable resolution.

## Methods

### Cross Tube Phantom experiments

To gauge the effectiveness of DSR we used a cross tube phantom^23–25^ comprising of two thin-walled cellulose capillary tubes (Hemophan®, Membrana) of internal diameter 200 μm ± 15 μm, a wall thickness of 8 ± 1 μm in the dry state, and a length change under wet conditions of ± 1%. We placed these two tubes close together in a cross formation, with cells pulled upwards by a syringe pump (Pump 33 DDS, Harvard Apparatus, Massachusetts, USA). These crossed tubes allow for the measurement of these two tubes are a wide range of distances from each other, allowing for discernment of the imaging resolution by observing the point at which the tubes become indistinguishable from each other. Imaging was conducted using an L11-4v probe and a Verasonics Vantage 256 system transmitting 3 cycle pulses with 4 MHz centre frequency. The cells we used were MDA-MB-231-mARG_Ana_ cells as described in Hurt *et. al*.^13^. In brief, we engineered MDA-MB-231 cells to constitutively co-express the plasmids shown in Figure 3 (top) which enable expression of GVs derived from anabaena, GFP, and BFP when doxycycline induced (1 mg/mL). We then purified these cells via fluorescent sorting for co-expression of GFP and BFP and grew them out over several passages.

### Buoyancy purification experiments

We buoyancy purified cells by adding density medium (OptiPrep Density Gradient Medium, SigmaAldrich, Missouri, USA) to the media to achieve final densities of 1.04 g/mL (pure) or 1.02 g/mL (purer), and then centrifuged (500 g, 5 minutes). To explore this further we split the additionally placed the purified samples into a flow cytometer (Attune NxT Flow Cytometer, Thermo Fisher Scientific, USA). Investigating the levels of production of GFP and BFP after gating out the cells using forward side scatter (FSC).

### Deactivation Imaging

We generated a series of six high pressure (6.2 MPa peak positive pressure) B-Mode images at 200 frames per second, 3 cycle pulses. In the first of these six pulses (here called the deactivation pulse) the protein shell of the GV is fractured^22^, releasing the gas within. This gas then cavitates over the remaining cycles, releasing a strong acoustic signal, after which the bubble collapses. On the subsequent pulses the GVs are no longer present, as such the signal that is reflected to the transducers for these pulses represent all the structures in the imaging field except for the GVs. By taking the median image intensity of the five images after the deactivation pulse and subtracting this from the deactivation pulse we end up with a deactivation image which contains the isolated signal from the GVs (Fig. 1).

### DSR pipeline

To super-localise the bubbles we implemented the processing pipeline shown in Figure 2. After generating the deactivation image, a noise threshold was applied. This was then followed by a more aggressive peak-detecting adaptive threshold as described by Bradley *et. al*.^28^ with a sensitivity of 0.1. With the noise successfully removed from the image we apply a normalized 2D cross-correlation with a 2D gaussian who’s width was empirically derived to match the size of the point spread function. The effect of this is to generate an image which contains maxima at the centre of the areas where the deactivation image bears the closest resemblance to the point spread function. We then threshold this correlated image, size gate it to ensure only points the size of a cell are considered, and use the centroids of the thresholded images to generate an estimate of the location of the super-resolved cell. For display purposes we then convolved with a 2D Gaussian (FWHM 0.15 mm) to produce the final SR image.

## ACKNOWLEDGMENTS

The authors gratefully acknowledge funding from the International Human Frontier Science Program Organization (grant LT0036/2022-L), The President’s PhD Scholarships (L. Taylor), the National Institute for Health Research i4i (Grant NIHR200972), the National Institutes of Health (R01NS120828 to M.G.S.), the BBSRC (BB/W001497/1), and the Chan Zuckerberg Initiative.

